# Particle localization using local gradients and its application to nanometer stabilization of a microscope

**DOI:** 10.1101/2021.11.11.468294

**Authors:** Anatolii V. Kashchuk, Oleksandr Perederiy, Chiara Caldini, Lucia Gardini, Francesco S. Pavone, Anatoliy M. Negriyko, Marco Capitanio

## Abstract

Accurate localization of single particles plays an increasingly important role in a range of biological techniques, including single molecule tracking and localization-based superresolution microscopy. Such techniques require fast and accurate particle localization algorithms as well as nanometer-scale stability of the microscope. Here, we present a universal method for three-dimensional localization of single labeled and unlabeled particles based on local gradient calculation of microscopy images. The method outperforms current techniques in high noise conditions, and it is capable of nanometer accuracy localization of nano- and micro-particles with sub-ms calculation time. By localizing a fixed particle as fiducial mark and running a feedback loop, we demonstrate its applicability for active drift correction in sensitive nanomechanical measurements such as optical trapping and superresolution imaging. A multiplatform open software package comprising a set of tools for local gradient calculation in brightfield and fluorescence microscopy is shared to the scientific community.

## 1. Introduction

Localization of micro- and nanoparticles has a broad applicability and plays a significant role in different physical and biological methods. Some examples include position and force measurement in optical tweezers [1,2], investigation of motility of swimming microorganisms [3], tracking of objects in microfluidic devices [4], study of dynamics of proteins and vesicles in cells [5,6], and superresolution fluorescence microscopy [7]. Some advanced imaging techniques, such as single molecule localization microscopy [8], heavily rely on accurate subpixel localization of fluorophores with numerous tools available for superresolution image reconstruction [9–11].

Another class of applications in which a position of particles needs to be determined quickly and precisely is the active mechanical stabilization in optical microscopes [12]. In experiments that require nanometer or subnanometer stability a feedback system is crucial. For such setups, the thermal drift of the viewing plane due to heating/cooling of the optical elements creates significant issues. This occurs, for example, in nanomechanical measurements performed with optical and magnetic tweezers or atomic force microscopes [13]. If the imaging system is combined with optical tweezers, thermal drifts become even more severe due to the presence of a high-power laser beam. A similar problem arises in single molecule localization microscopy (SMLM) as methods like STORM (STochastic Optical Reconstruction Microscopy) and PALM (Photo-Activated Localization Microscopy) [14] require high power of the excitation beam and long acquisition times and, therefore, will greatly suffer from an imaging plane drift. While there are algorithmic solutions to correct for this displacement after image acquisition, it is not always feasible as the drift might be too large to be compensated in a postprocessing. Moreover, this approach is not applicable to nanomechanical measurements. An active stabilization system that controls the position of the objective or sample chamber can compensate for the drift in the recorded data in the first place. Most commonly such systems utilize a bead or fluorescent marker attached to a coverslip as a reference for correction and require the position of the particle to be measured in three dimensions. The most common way to localize a single particle is to apply a threshold to select the brightest pixels in the image followed by calculation of an intensity weighted centroid. Despite being very fast, this method shows poor performance and has several practical issues [15]. The aforementioned tracking algorithms in superresolution fluorescence microscopy were developed for a specific task of image reconstruction for fluorescent probes and are unsuitable or poorly applicable in other cases. Another interesting approach was proposed in [16]. The DeepTrack software utilizes recent advances in convolutional neural networks to localize particles of different types, shapes and sizes. However, as with all artificial neural networks, it requires a training data to operate and its performance heavily relies on the size and quality of the dataset provided.

Here, we present methods for particle localization and microscope stabilization based on the calculation of local gradients of the image intensity. In general, the local gradient algorithm (LoG) can be useful in different applications in image processing that require calculation of gradients. However, for the purpose of this paper, we will focus solely on the use of local gradients in particle localization tasks. We propose a set of tools for 3D localization of both fluorescent and unlabeled particles. The software was primary developed for active stabilization systems in sensitive biological experiments and, therefore, includes parameters which can be easily adapted to a specific conditions, executes in a short time-frame to allow high processing rates, provides accurate results at low signal-to-noise ratios. However, the LoG algorithms presented here have a much more broad applicability. We test and demonstrate usability of the LoG algorithm in XYZ-localization of particles in brightfield imaging and fluorescent particles in astigmatism based microscopy [17].

To make the software more suitable for immediate incorporation the LoG tools are available on the platforms which are commonly used for data capture and analysis: Matlab, LabVIEW and Python. The software can be obtained from [18,19].

## 2. Results

We define a local gradient in a given point as the intensity weighted centroid of all the pixels within a radius *r* from that point (see Fig. 1). By calculating local gradients for each pixel we obtain horizontal and vertical gradient matrices (*G*_*x*_ and *G*_*y*_ on Fig. 1) of the original image (see Supplementary S1 section for more detailed derivation).

**Fig. 1.**
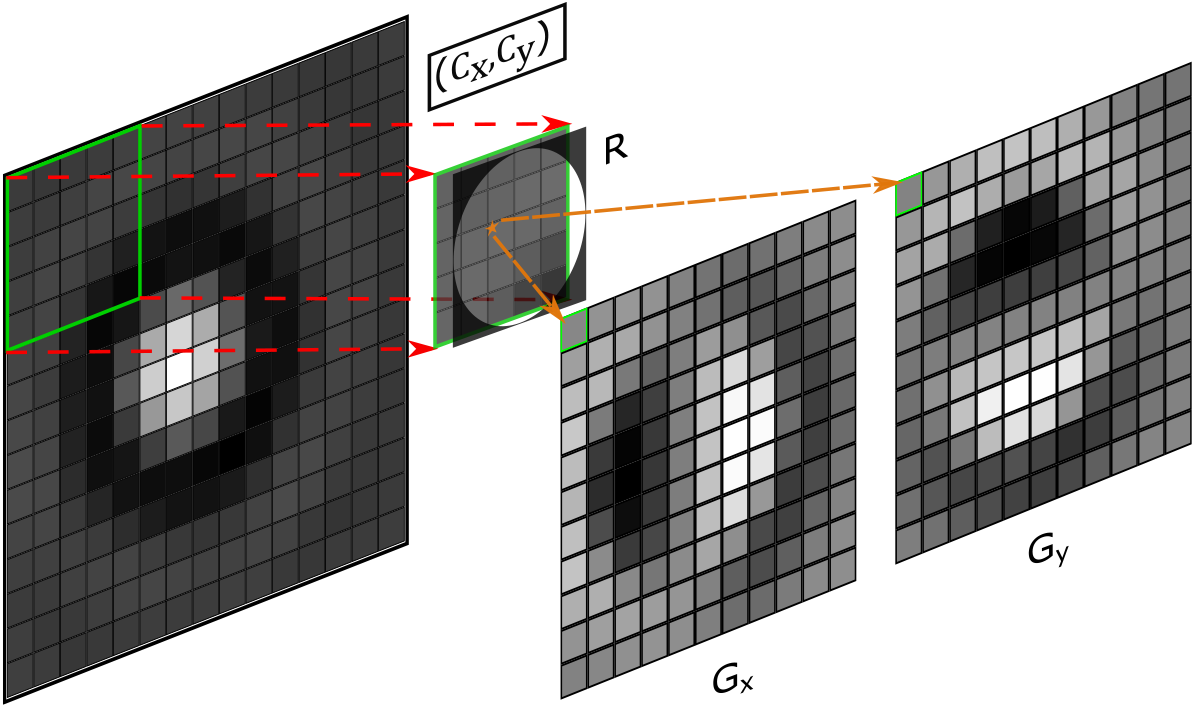
Visualization of the local gradient algorithm. For an *m*×*n* image (here 15×15) a centroid of a sliding window size *K* = 2⌈*r* − 0.5⌉ + 1 (here 5×5 with *r* = 2.5) is calculated. Each pixel of gradients *G*_*x*_ and *G*_*y*_ is determined as the x (*C*_*x*_) and y (*C*_*y*_) coordinates of the centroid correspondingly, which is calculated relative to the center of the window. The resulting matrices *G*_*x*_ and *G*_*y*_ have the size (*m* − *K*) × (*n* − *K*). Negative gradient values for images *G*_*x*_, *G*_*y*_ are represented as darker pixels and positive as a whiter pixels. An orange star depicts the centroid. *R* is a circular mask *K* × *K* of radius *r*.

The use of gradients to localize radially symmetrical single particles has been shown to be a robust and effective method [20,21]. Though such methods demonstrate high accuracy and stability to brightness variations, they are susceptible to noise and show poor performance at low signal to noise ratios. Also most of them lack flexibility due to employment of a fixed-size kernel for gradient calculation. In the proposed LoG algorithm the radius of the window *r* determines the number of pixels included in the calculation of local gradients. Therefore, by adjusting *r* one can enhance a calculated gradient for an object of a specific size. Moreover, this provides much better results for images with low SNR as more pixels will averaged.

We show and thoroughly test several approaches based on LoG algorithms to determine 3D position of particles in both brightfield and fluorescent microscopy.

### 2.1. xyz position detection in brightfield microscopy

Figure 2 a-c depicts the localization of a 0.9*µm* silica particle from its brightfield image (Fig. 2a). The gradient matrices *G*_*x*_ and *G*_*y*_ calculated from the LoG algorithm using Eq. S3,S4 form a vector field (as depicted by blue arrows in Fig. 2c) that contains a gradient vector for each pixel. The center of a radially symmetrical particle can be determined as an intersection of all gradient lines. However, the presence of the noise or uneven illumination in the background of the image will create gradient vectors with random or incorrect orientation that may disrupt the estimation of the center. To limit the influence of such artifacts on the localization the magnitude (Euclidean norm) of gradient vectors is used to exclude low-magnitude values (Fig. 2b,c). The calculation of an intersection point of gradient lines implies solving a system of linear equations. In practice such system is overdetermined and inconsistent as it is very unlikely that all the gradient lines will intersect at a single point. Therefore, the center of the particle is calculated using the method of least-squares applied to gradient lines with the highest magnitude of gradient vectors (see Supplementary S2).

**Fig. 2.**
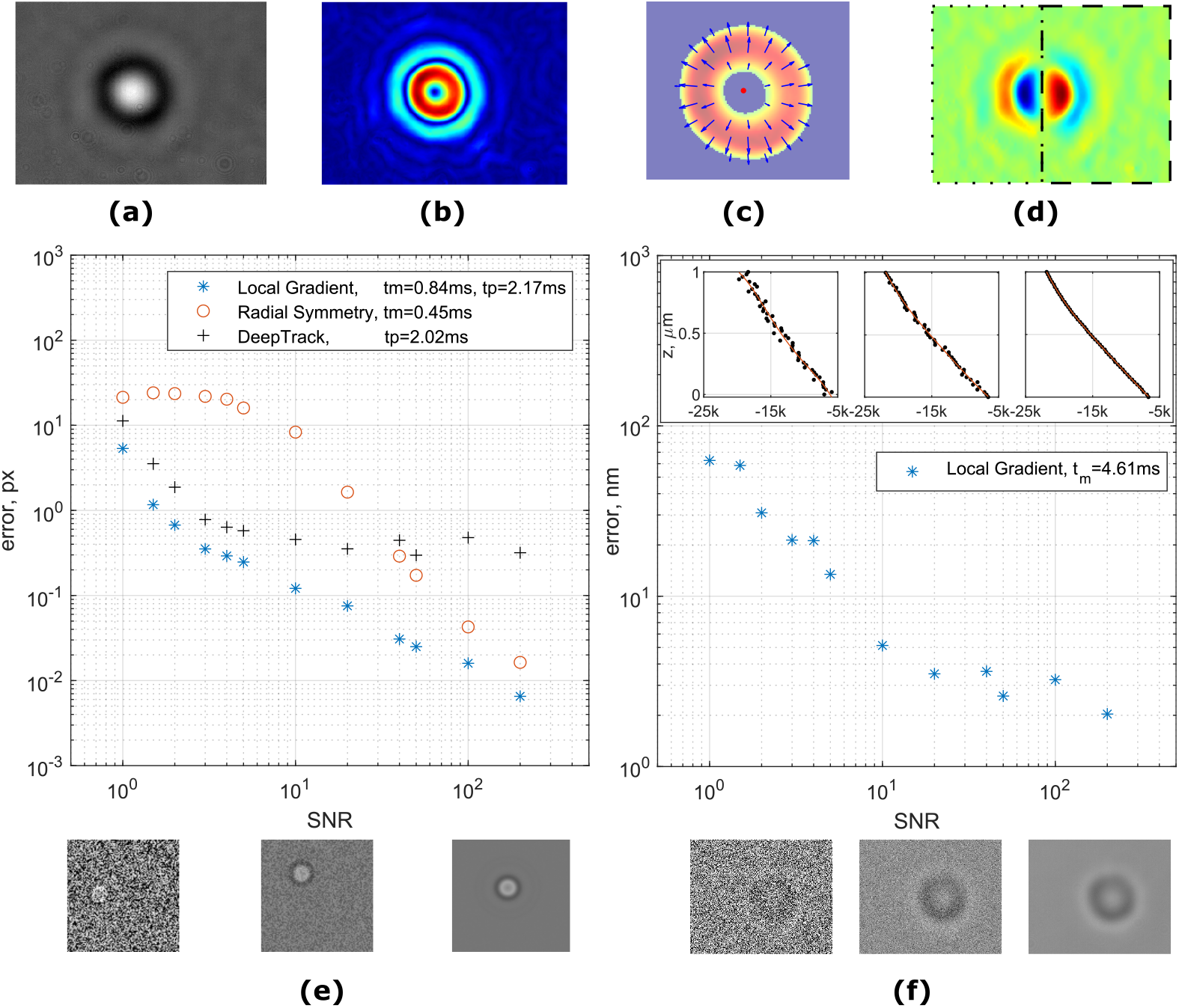
Localization of a particle using local gradient algorithm (images a-d are not to scale for better visualization). (a) Input image of the 0.9*µm* silica particle. (b) Magnitude of local gradients. (c) Magnitude of local gradients after thresholding. Arrows show the direction of gradients (from high to low). The position of the particle (red spot) is determined as a least-square intersection of all gradient vectors. (d) z-value calculation. Local gradient *G*_*x*_ is split into left (dotted) and right (dashed) parts. z-value is determined as a difference between the sums of gradient values of “right” and “left” sides. (e) Accuracy and execution time performance of XY-localisation algorithms and different SNRs. Each error point is an average error of 20 images. tm and tp are average execution times in Matlab and Python correspondingly. Example of the test images is shown under the plot. (f) Accuracy and execution time of LoG algorithm for z-localisation. Inset shows examples of calibration curves.

To investigate the performance of the LoG algorithm for xy-localization we run a set of tests and compare the results to other available methods: radial symmetry algorithm [20] and DeepTrack software [16]. The first method uses a similar approach in the calculation of the center of the particle – least-square intersection of gradient lines, but significantly differs in the gradient calculation. In the radial symmetry algorithm the gradients are calculated between adjacent pixels only, which leads to increase in the detection error due to noise amplification for images with low SNR. The second software package, DeepTrack, provides tools for training and validation of a convolutional neural network (CNN) for particle tracking. To compare our results with DeepTrack we have trained a network using built-in tools feeding 30000 images of simulated particles randomly positioned and with added noise (see Methods 4.3). It should be noted that the performance of the DeepTrack network, as in any CNN, strongly depends on the type and size of the training data. Thus, the test results for a given network will serve as a reference of what is available out of the box and not necessarily reflect the full capability of the software. As DeepTrack is based on Python, and radial symmetry algorithm is written in Matlab, we compare each of the software packages with a corresponding implementation of the LoG algorithm.

The test results are shown in Fig. 2e. The LoG algorithm demonstrates an excellent noise stability and outperforms both radial symmetry and DeepTrack in most cases. The execution time of all methods is comparable for a given platform. A precalculation of the Fourier transform of matrices in Eq. S5 saves 0.30*ms* (∼24%) which may speed up the execution time in case of using the same parameters for multiple images.

The axial position of the particle in brightfield microscopy is a more complex task. Using local gradients we have devised a simple yet fast and effective approach to measure z-position. The basic concept is shown in Fig. 2d. A horizontal local gradient image *G*_*x*_ is split into two parts relative to the center of the particle. The difference between the sum of all gradient values on the “right” and “left” sides provides an excellent metric for calibration curve or look-up table. A similar metric can be build for the vertical gradient *G*_*y*_ and an average of both horizontal and vertical gradients constitutes a z-position value. The test results of the noise performance shown in Fig. 2f were performed using real images of a single particle with artificially added noise. A set of 101 images is taken at different heights with step size of 10nm. In order to differentiate an input data, each odd image was taken to create a calibration curve while each even image is used for the test (see Methods 4.1).

As there are no “pixels” in the axial direction, we conclude the test results basing on the nanometer position of the objective piezo scanner. With such a simple approach we are capable of achieving an error as low as ≈ 2*nm* in the case of high SNR images.

### 2.2. xyz-localisation in astigmatism-based fluorescence microscopy

Determination of the position of fluorescent particles and markers in *x* and *y* is a very similar task to the localization of particles in brightfield microscopy. Hence, the same methods can be applied. The difference between the two is that fluorescent images, in particular when a sample is at a single molecule concentration, are more noisy due to a typically low signal from fluorophores. Figure 3d demonstrates a comparison of accuracy of local gradient method, radial symmetry and DeepTrack for generated gaussian-like fluorescent particles (see Methods 4.2).

**Fig. 3.**
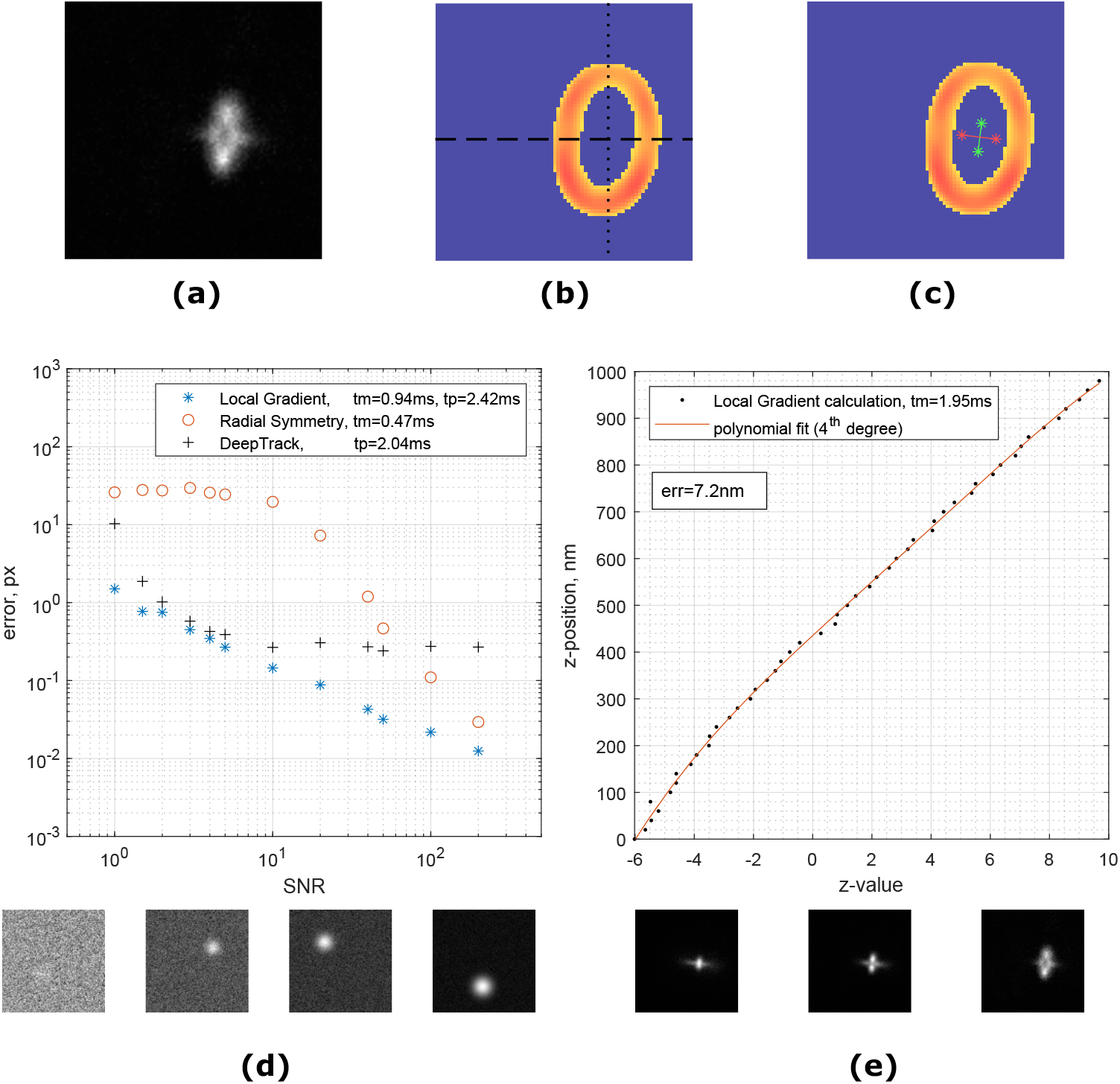
(a) Image of a single fluorescent particle (polystyrene, 0.51*µ*m) attached to a coverslip. Astigmatism is introduced by a cylindrical lens and the imaging plane is ≈500*nm* above the surface (b) Magnitude of local gradients. Dashed and dotted lines are showing the top/bottom and left/right split of the local gradient images for z-value estimation correspondingly. (c) Two axes (green and red lines) are built from the centers of splitted gradient lines (d) Comparison of algorithms for localization of a simulated gaussian-like particle at different noise levels. tm and tp are average execution times in Matlab and Python correspondingly. Examples of the test images are shown under the plot. (e) z-value calibration curve in astigmatism based microscopy. The average error for predicting a z-position of the particle is 7.2*nm*. Examples of the test images are shown under the plot.

Measurement of the axial position in case of fluorescent particles is much more challenging. Unlike the image of micron-sized beads in brightfield microscopy, the point spread function (PSF) of fluorescent particles with subdiffraction dimension does not vary significantly in shape at different heights near the objective focal plane. This causes the particles below and above the imaging plane look the same and makes them very hard to properly localize. The solution to this issue is to defocus the image [22,23] or introduce some sort of aberrations into the imaging system to break the axial symmetry of the PSF [24,25]. One of the methods to provide sensitivity in axial position for fluorescent probes is to use a cylindrical lens in the optical path [17]. The PSF remains roughly round only when the particle is in the imaging plane and becomes elliptical when outside with major axis flip of 90 degrees when passing through the imaging plane (see Figure 3e). It has been shown that in such system the z-position is proportional to the ellipticity of the PSF. The LoG algorithms provide an opportunity to utilize their features to create a metric for fast and efficient axial localization. A local gradient image of the fluorescent particle with introduced astigmatism can be easily made to resemble an ellipse by adjusting the window size *r* (Figure 3b).

The calculation procedure of the axial metric is shown in Figure 3b,c. The image is thresholded by the magnitude of the gradient. Once the xy-position has been found, the image is split into top/bottom or left/right sides relative to the center of the particle. For each part a least-square intersection of gradient lines is calculated resulting in four points which form two axes. The length of the major axes is taken as a z-value (see Supplementary S4 for detailed description).

The results of the noise tests for xy-localization (Figure 3d) show that, again, the LoG algorithm outperforms both DeepTrack and Radial Symmetry at all tested SNRs while keeping the execution time comparable to other methods. For z-position measurements the proposed z-value metric provides a linear response within the tested 1000*nm* displacement with average execution time of 1.96*ms* per image. The averaged error in position determination is *z*_*err*_ = 7.2*nm*

### 2.3. Feedback in brightfield microscopy

Finally, the xyz algorithms for brightfield images described above were applied for a feedback system to stabilize the drift of the imaging plane and reduce the impact of mechanical noise. The experimental setup was an ultrafast force-clamp spectroscopy system [26], which is used to study protein interactions under a constant load. In these experiments, a “dumbbell” structure consisting of an actin [27–29], microtuble [30], or DNA filament [31] strained between two microbeads is held by optical tweezers. The actin filament is brought into vicinity of a stationary microbead that is attached to the coverslip and covered with proteins of interest. As the interaction area between the filament and the protein is often on the order of nanometers a mechanical stabilization is critical for protein attachment/detachment to be observed and measured. Moreover, measurement of nanometer-sized conformational changes of proteins critically relies on the mechanical stability of the system. Therefore, in these experiments the stationary microbead is used as a fiducial mark for microscope stabilization [32]. Here, we demonstrate the use of local gradients for the mechanical feedback system in stabilization of the viewing plane (see Methods 4.4). Figure 4a shows tracking of a single particle attached to the coverslip with and without feedback.

**Fig. 4.**
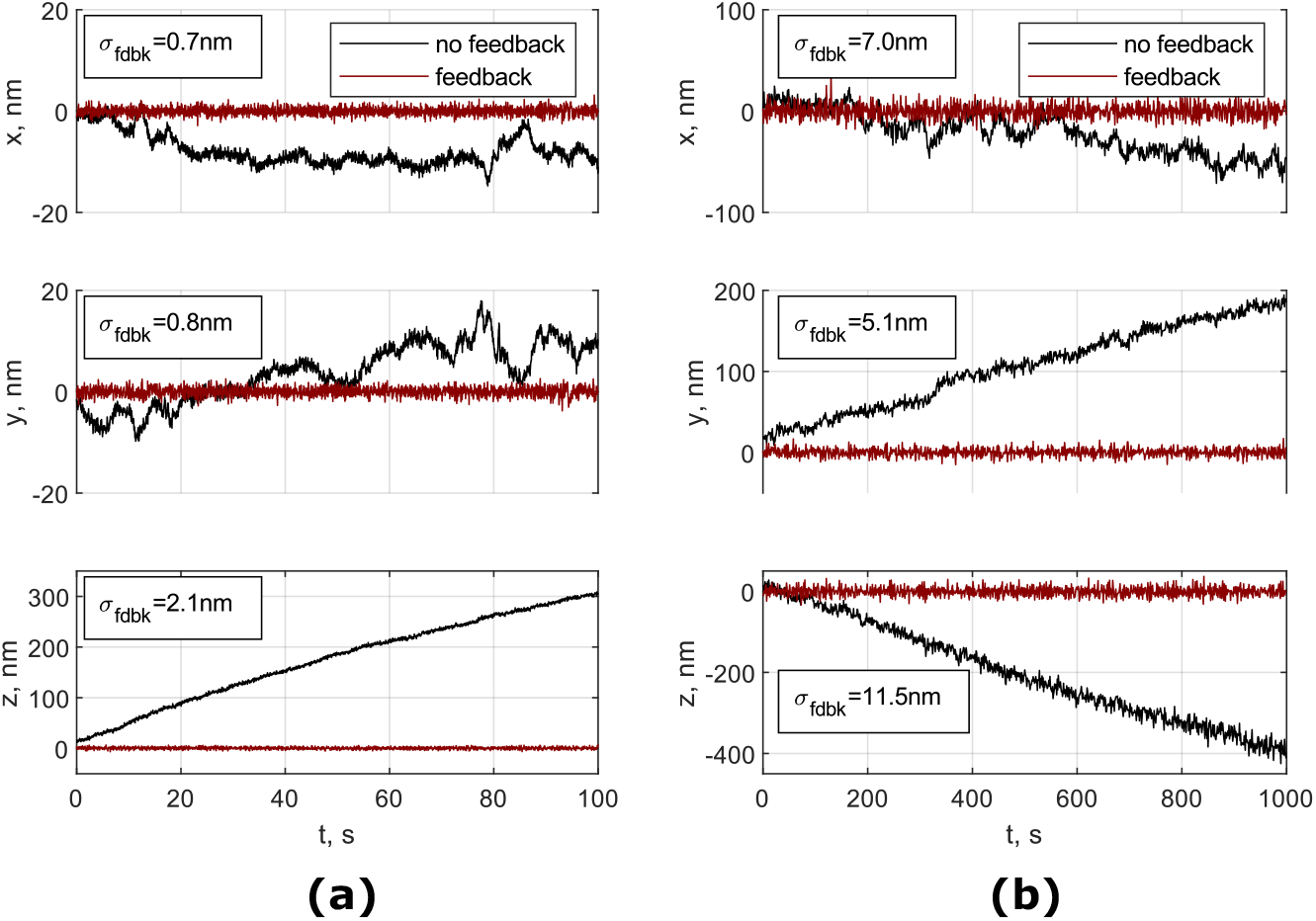
3-D tracking of a spherical silica particle in brightfield microscopy (a) and fluorescent polystyrene particle in astigmatism microscopy (b) that were attached to the coverslip with feedback system on and off. Inset indicates a standard deviation of the signal with feedback on.

Our feedback system demonstrates ability to suppress mechanical noise and thermal drifts to sub-nanometer levels in xy-localization and to 2*nm* in axial positioning. As can be seen from the plots, the displacement of the viewing plane with no feedback creates a sufficient movement, on the scale of tens to hundreds nanometers, which makes protein interaction measurements in force-clamp setups impossible to do

### 2.4. Feedback in astigmatism-based fluorescence microscopy

A similar approach was used to test the performance of the feedback system using the LoG algorithm on fluorescent particles. This test was performed on an inverted fluorescence microscope for superresolution microscopy (see Methods 4.4) on a sample made of commercial 500 nm fluorescent polystyrene beads (Exc/Em 480/520) attached to a glass coverslip (see Methods 4.5). A single particle was tracked in 3-D for 1000 seconds (a representative time for STORM image acquisition). The calibration curve for z-value was recorded on the same bead before the acquisition. The results for both feedback controlled and free-running cases are shown on Fig. 4b. Again, we were able to demonstrate a stable positioning of the sample with a standard deviation of the position in the range of 5 −7*nm* for x-y localization and 11.5*nm* for z localization.

Next, we applied the feedback system to record a 3D-STORM image of the actin cytoskeleton of a mammalian cell using a fluorescent bead as fiducial marker. A 3D-STORM image of the actin cytoskeleton of a different cell from the same sample was recorded without the feedback for comparison (Fig. 5a).

**Fig. 5.**
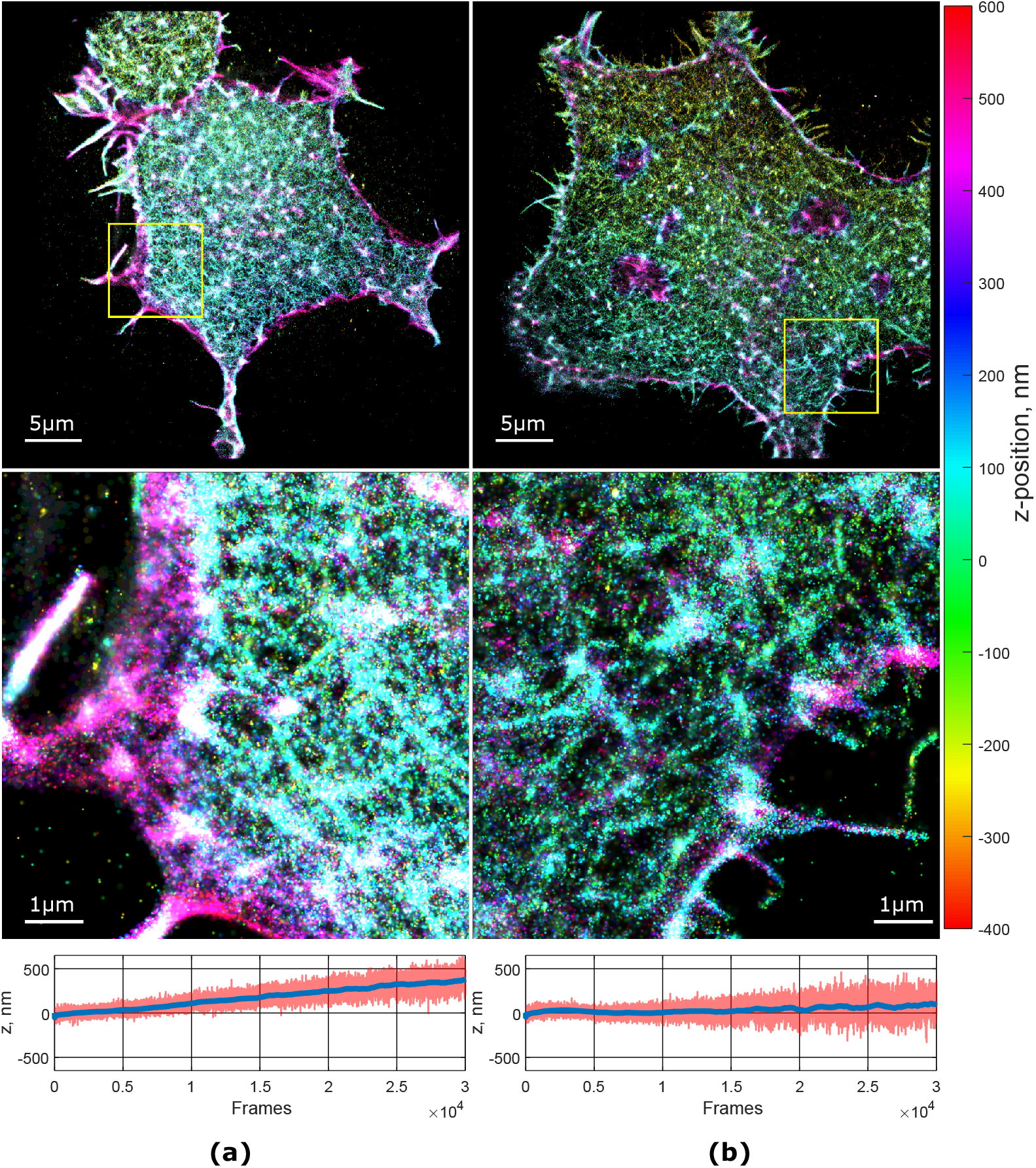
Reconstructed 3D STORM images of cells without feedback correction (left) and with feedback (right).Plots under the images show the change in the average z-position (red line) of all detected fluorophores (blue line represents moving average of 1000 points).

Additionally, we estimate the drift in the z-axis by calculating the average position of all detected fluorophores for each frame. The actin cytoskeleton has an uneven distribution within the cell volume, with a dense branched cortex spanning about 200-300 nanometers from the cell membrane and a less dense network protruding towards the cell nucleus. If the viewing plane is drifting axially the average z-position of all fluorophores in a frame will drift as well. The axial drift is clearly visible in the images acquired in the absence of feedback, contrary to what is observed with the feedback (plots in Fig. 5). As a consequence, the image acquired with the feedback (Fig. 5 on the right) appears sharper and shows more fine details compared to the one without feedback.

## 3. Discussion

We have developed and tested several methods for particle localization utilizing local gradients. There are multiple advantages in the use of LoG-based algorithms. Low dependence of the gradient images on the background intensity level makes them suitable for particle detection in changing conditions or uneven illumination. A variability of the window size *r* allows selective enhancement of the objects of particular size (see Supplementary S6). Though the particles should significantly differ to be selected, this feature helps removing small dust spots from the image or filter particles by size in strongly polydisperse samples. Also, as LoG xy-localisation algorithms are based on symmetry of particles, they can be applied to all kind of objects with radial symmetry regardless of their origins (fluorescent, brightfield or holographic imaging). Furthermore, these methods are applicable to particles which are only partially present in the field of view of the camera as even a small number of gradient lines will point towards the center. This property can be useful in case of limited field of view or obstruction of the particle by other objects.

In comparison with other methods, LoG algorithms are more noise resistant and show better accuracy for all tested cases. While Deeptrack can be trained to track much broader classes of objects, LoG software is more flexible and provides better control over the tracking process for radially symmetrical particles. For mechanical stabilization systems the execution time is an important parameter for the localization software to reach sufficient feedback framerate. In LoG methods the execution time mostly depends on the input image size as the most time-consuming step are convolutions represented as a series of forward and inverse Fourier transforms. It should be noted that the speed of FFT algorithms used for Fourier transform calculations has a complex dependence on the image size and can be optimized by cutting/expanding the input image to the appropriate size.

For accurate determination of the axial position a calibration curve or look-up table should be created for each particle of interest as small variations in size may lead to much higher errors than those obtained in the tests. However, while the method is less accurate than the xy-position algorithm, it shows an excellent precision. For a mechanical feedback system the accuracy is usually of less importance than the precision as the required outcome is stability of the viewing plane at some relative position. This makes it possible to apply a prerecorded calibration to similar particles. The main requirement for the metric in this case is a monotonic output across the possible axial ranges, which our algorithms immaculately fulfill even when the viewing plane is crossing the center of the particle.

The LoG algorithms were primarily developed for feedback systems that require fast and precise determination of the 3D position of the particle. We have demonstrated an efficient sub-nanometer stabilization of a microscope stage that is used for force spectroscopy of proteins. It is clear that a free-running system will not allow any protein interaction measurements due to a significant drift which will quickly move the proteins out of interaction region. A similar problem exists in superresolution microscopy in which a stack of images is recorded (such as PALM/STORM). In many cases the movement of the viewing plane can be corrected in post-processing by drift-correcting algorithms. These algorithms are especially effective in xy-axes but work only in a limited range of axial drift as the SNR of out-of-focus chromophores quickly decreases making them undetectable. We have shown that a feedback system based on local gradient algorithm shows significant improvement in z-axis stability and improves the overall image quality.

## 4. Methods

All calculations were performed on Lenovo ThinkPad T15g Gen 1 Core i7-10850H vPro, Windows 10 Pro 64, 32 GB (2933 MHz).

### 4.1. XYZ detection accuracy tests

Images of particles for xy-performance tests in brightfield were obtained from a radial profile of a 3*µm* silica bead (Bangs Laboratories, SS05001, M.D. 3.17*µm*) fixed on the coverslip. A smoothing spline is applied to reduce a pixelation in the image. A set of 20 images (100 × 100pxls) was generated for each SNR (12 SNR levels) with randomly positioned particles. The same set of images is used for all the software packages to test. The error is defined as a distance between the predicted and true location of the particle. Local gradient parameters used to track the particles are: *r* = 10, threshold cut-off 1/1.4 of maximum gradient intensity. The execution time was estimated by performing 200 runs of 240 generated images (48000 runs in total). To equalize a cross-platform productivity, Tensorflow was forced to use a single core.

Axial positioning is tested on set of images (222*x*276 pxls) of 1*µm* silica particle (Bangs Laboratories, SS0400, M.D. 1.05*µm*) attached to the coverslip. The image of the particle is recorded at different heights with 10 nm step using objective scanner. The calibration is created by using every odd image (20nm step, 51 images) and the tests are performed on each even image (20 nm step, 50 images). A random noise was added at specified SNR (12 levels). Local gradient parameters used to track the particles are: *r* = 35, threshold cut-off 1/1.5 of maximum gradient intensity.

### 4.2. XYZ detection for fluorescent particles

Images for the x-y positioning tests (100×100 pxls) were generated from a high-resolution images (10000×10000 pxls) of a randomly distributed 2D gaussian function. A set of 20 images is generated for each SNR (12 SNR levels) with randomly positioned particles. Local gradient parameters used to track the particles are: *r* = 12, cut-off threshold 1/1.3 of maximum gradient intensity. The execution time was estimated by performing 200 runs of 240 generated images (48000 runs in total). Tensorflow was forced to use a single core.

Axial positioning is tested on a set of images of 500*nm* fluorescent beads (100×100 pxls) attached to the coverslip. The image of the particle is recorded at different heights with 10 nm step using objective scanner. The calibration is created using every odd image (20nm step, 51 images) and the tests are performed on each even image (20 nm step, 50 images). Local gradient parameters used to track the particles are: *r* = 10, cut-off threshold 1/2 of maximum gradient intensity.

### 4.3. Training of the DeepTrack CNN

The Deeptrack software [16] was used to create a convolutional neural network for particle detection. Along with the amount, quality, and variety of the data used to train the neural network, the architecture of the neural network, its training parameters and conditions are key factors in obtaining a good result. Here we build and train the network similarly to the examples and tutorials shipped with the software package.

The neural network model consists of four convolutional layers with the number of output filters equal to 16, 32, 64, 128 respectively and two subsequent dense layers with sizes of 64 neurons each. Nonlinear activation functions for dense layers are set to Rectified Linear Unit (ReLU). Each convolutional layer had 0.2 dropout, valid pooling block, and steps per pooling equal to 1. Adam optimizer was applied. Mean squarederror was used as a loss function and pixel error as a metric. The custom pixel error function: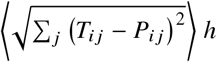, where *i* = 2, *j* is a batch size, *h* is an image size, *T*_*ij*_ and *P*_*ij*_ are the true and predicted coordinates of the particle correspondingly. The following parameters are used within the DeepTrack software to generate test images: Scatterer: PointParticle, Optics: Fluorescent microscope (NA=0.7, wavelength=660 nm, resolution=1e-6, magnification=25, refractive index medium=1.33, upscale=2, padding=30); Since the size of test images is larger than the size used in the examples (100×100 versus 64×64), one more convolutional layer was added and the number of neurons in dense layers was doubled. Neural networks were trained on a dataset of 20000 images verified at 512 validation images. The batch size was equal to 64. The number of training epochs was set to 250 epochs with early stopping equal to 20 epochs. By setting an early stopping parameter, Keras was able to stop training in case of a stop of the loss function decrease. In reality, the number of training epochs did not exceed one hundred.

Background and random Poisson noise were added to bring images closer to the real experimental data. (background =1, Poisson noise min/max SNR 2/80. All generated images were augmented and had the size of 100 ×100 pixels. The output of the network is two parameters - x and y coordinates of the particle center.

### 4.4. Feedback

The mechanical stabilization in brightfield microscopy was implemented on an existing ultra-fast force-clamp system. The setup includes a custom-built microscope with piezo controlled stage (Physik Instrumente, P-527.2CL) and PIFOC Objective Scanner (Physik Instrumente, P-725.4CL). To reduce ambient noises the system is built on a table resting on a supports with pneumatic noise suppressors. Additionally, the microscope is placed on top of elastomeric dampers. The particles were imaged with water immersion objective (Nikon Plan-Apo 60X, N.A. 1.20) onto a USB CMOS Camera (Thorlabs DCC1545M). The pixel-to-nm ratio of the camera was 18.3*nm*/ *pxl*. z-value calibration was obtained prior to the feedback test on the same particle by scanning the objective within ±500*nm* range (100*nm* step size) followed by a linear fit of the calculated z-values.

3D STORM imaging was performed on an inverted wide-field fluorescence microscope (Nikon ECLIPSE TE300) with 643 nm and 488 nm excitation lasers. Excitation was performed with inclined illumination through a TIRF 60 x objective (Nikon 60 X, oil immersion, NA 1.49 TIRF) to optimize the image contrast. Emitted fluorescence was collected through the same objective and imaged on an EMCCD Camera (Andor iXon X3) after an additional 3x magnification. Full field of view is 40×40 *µm*^2^ wide, with 80 *nm* pixel size.

### 4.5. Fluorescent microspheres sample preparation

Dragon Green beads (Bangslab, FSDG003, 0.51 µm) were diluted at 0.1 % (v/v) in 25 mM MOPS, 25 mM KCl, 4 mM MgCl2, 1 mM EGTA, 1mM DTT pH 7.2. A chamber, composed by a standard glass coverslip sandwiched on a microscope slide with double-sided sticky tape, was filled with the beads solution and incubated for 5 minutes at room temperature. After a careful wash to remove unattached beads, the chamber was sealed with silicon grease and put on the microscope stage for measurements.

### 4.6. STORM imaging

STORM sample consists of fixed HEK 293T cells with Dragon Green microspheres (Bangslab, FSDG003, 0.51 *µm*) as reference beads for the feedback algorithm. Cells were plated on poly-L-lysine coated 18-mm diameter glass coverslips. After 24 hours incubation at 37 °C cells were fixed, by incubating with 4% paraformaldehyde (PFA) solution for 10 minutes, permeabilized in 0.075% Triton X-100 solution for 7 minutes and blocked with 4% bovine serum albumin (BSA) solution in PBS with added Ca++ Mg++ for 30 minutes. After blocking, a dilution of Dragon Green beads at 0.1% (v/v) in 25 mM MOPS, 25 mM KCl, 4 mM MgCl2, 1 mM EGTA, 1mM DTT pH 7.2 was incubated with cells on the coverslip for 2 minutes. Then the actin cytoskeleton was labelled by incubating overnight with 0.5 *µ*M Alexa Fluor 647 phalloidin at 4 °C. After 20 hours, the sample was washed once with PBS and mounted on an imaging chamber with imaging buffer composed by 200 mM *β*-MercaptoEthylamine hydrochloride (MEA), 20% (v/v) sodium DL-Lactate solution, 3% (v/v) of OxyFluor, in PBS pH 8. Prior to final rendering of the super-resolved images localizations with a lateral uncertainty greater than 150 nm were filtered out. Final images were visualized at 10x magnification (i.e. the image pixel size is 8 nm). Images acquired with 3D STORM were reconstructed with ImageJ plugin ThunderSTORM. The details of the reconstruction parameters can be found in the Supplementary S5.

## 5. Acknowledgements

This work was supported by the European Union’s Horizon2020 research and innovation program under grant agreement no 871124 Laserlab-Europe. A.V.K. was supported by the Human Frontier Science Program Cross-Disciplinary Fellowship LT008/2020-C. O.P. and A.M.N. were supported by the project 1.4. B/185 National Academy of Sciences of Ukraine.

## 6. Contributions

A.V.K. conceived the project, developed and tested algorithms, wrote the code in Labview and Matlab. O.P. and A.M.N. wrote and tested Python version of the code and performed training and tests with Deeptrack. A.V.K., L.G. and M.C. designed test experiments. A.V.K. performed brightfield microscopy feedback tests. C.C., F.S.P., L.G. and A.V.K. performed superresolution imaging tests. M.C. provided general supervision of the project. A.V.K. wrote the manuscript with input from all authors.

