## Supplementary material for "Particle localization using local gradients and its application to nanometer stabilization of a microscope"

#### *S1. Local gradient algorithm*

For a given image $I(x, y)$, we define a local gradient of a pixel $(x_i, y_j)$ as a centroid of all pixels within a circle of radius $r$ $\{r \in \mathbb{R} \mid r > 0.5\}$ centered at the $(x_i, y_j)$:

$$\left(g_x^{(i,j)}, g_y^{(i,j)}\right) = \left( \frac{\sum\limits_{k,l=-r}^{r} I\left(x_{i+k}, y_{j+l}\right) x_{i+k}}{\sum\limits_{k,l=-r}^{r} I\left(x_{i+k}, y_{j+l}\right)}, \frac{\sum\limits_{k,l=-r}^{r} I\left(x_{i+k}, y_{j+l}\right) y_{j+l}}{\sum\limits_{k,l=-r}^{r} I\left(x_{i+k}, y_{j+l}\right)} \right) \tag{S1}$$

By calculating gradients at each pixel the final gradient matrices are (also see Fig. 1):

$$G_{x,y} = \begin{pmatrix} g_{x,y}^{1,1} & g_{x,y}^{1,2} & \dots & g_{x,y}^{1,n} \\ \vdots & \vdots & \ddots & \vdots \\ g_{x,y}^{m,1} & g_{x,y}^{m,2} & \dots & g_{x,y}^{m,n} \end{pmatrix} \tag{S2}$$

Matrices $G_x$ and $G_y$ are $x$- and $y$-gradients correspondingly.

In most cases the calculation speed of the gradient can be increased by presenting the equations S1 and S2 as convolutions and applying the convolution theorem:

$$G_x = (I * X) \oslash S = \mathcal{F}^{-1}\left\{\mathcal{F}\{I\} \circ \mathcal{F}\{X\}\right\} \oslash S \tag{S3}$$

$$G_y = (I * Y) \oslash S = \mathcal{F}^{-1}\left\{\mathcal{F}\{I\} \circ \mathcal{F}\{Y\}\right\} \oslash S \tag{S4}$$

where "$\circ$" denotes Hadamard (element-wise) multiplication, "$\oslash$" denotes Hadamard division, $\mathcal{F}/\mathcal{F}^{-1}$ is a Fourier/inverse Fourier transform, "$*$" denotes convolution, $X$ and $Y$ are square matrices of size $k = 2\lceil r - 0.5 \rceil + 1$:

$$(X, Y) = \left( \begin{pmatrix} -r & -r+1 & \dots & r \\ -r & -r+1 & \dots & r \\ \vdots & \vdots & \ddots & \vdots \\ -r & -r+1 & \dots & r \end{pmatrix} \circ R, \begin{pmatrix} -r & -r & \dots & -r \\ -r+1 & -r+1 & \dots & -r+1 \\ \vdots & \vdots & \ddots & \vdots \\ r & r & \dots & r \end{pmatrix} \circ R \right) \tag{S5}$$

here $R$ is a circular amplitude mask of the same size as $X$ and $Y$. $S$ is the sum of all elements in each sub-image:

$$S = I(x, y) * J = \mathcal{F}^{-1}\{\mathcal{F}\{I\} \circ F\{J\}\} \tag{S6}$$

here $J$ is a square matrix of ones of size $k$.

For most tracking applications the size of the window will remain constant and therefore, $\mathcal{F}\{X\}$, $\mathcal{F}\{Y\}$ and $\mathcal{F}\{J\}$ can be precalculated and reused to reduce the total calculation time.

### S2. x-y position detection

The x-y position of the particle is determined as an intersection of all gradient vectors in a least-square sense. A system of linear equations to find the least-square intersection of gradient lines $\boldsymbol{p}$ is:

$$\boldsymbol{L}\boldsymbol{p} = \boldsymbol{q} \tag{S7}$$

$$\boldsymbol{L} = \sum_{j=1}^{K} w_j \left(\boldsymbol{I} - \boldsymbol{n}_j \boldsymbol{n}_j^T\right), \boldsymbol{q} = \sum_{j=1}^{K} w_j \left(\boldsymbol{I} - \boldsymbol{n}_j \boldsymbol{n}_j^T\right) \boldsymbol{a}_j \tag{S8}$$

where $w$ is a weighing vector, $\boldsymbol{I}$ is an identity matrix, $\boldsymbol{n}$ is a gradient direction vector, $\boldsymbol{a}$ is a coordinate vector of a point on the gradient line, $K$ is a number of lines/equations. The solution of the system of linear equations (S7) returns the center of the particle.

### S3. z-calibration curve

Figure S6 shows a calibration curve for $3\mu m$ particle.

### S4. z position detection in astigmatism based microscopy

Firstly, the x and y positions of the particle are calculated according to the previous section using the equations S3, S4 and S7. Then, a thresholded magnitude of local gradients is splitted vertically and horizontally relative to the calculated center. The gradient lines within each half are used in equation S7 to locate the position they are pointing to in a least-square sense. This results in four points which can be connected in two crossing axes similar to the minor and major axes in ellipse (however, in our case the axes may not be orthogonal). z-value is set as the length of the major axis with the sign determined as:

$$\begin{aligned} &\operatorname{sgn}(\sin(2\phi)(x_1 - x_3) + \cos(2\phi)(y_1 - y_3)) - \\ &\qquad - \cos(2(\phi + \pi/2))(x_4 - x_2) + \sin(2(\phi + \pi/2))(y_4 - y_2)) \end{aligned} \tag{S9}$$

where $\phi$ is the angle of the major axis which corresponds to positive z-values, $x_1 ... x_4$, $y_1 ... y_4$ are coordinates of the calculated centers for each half.

### S5. Thunderstorm parameters for analysis and visualization

Images acquired with 3D STORM were reconstructed through ImageJ plugin ThunderSTORM using the following parameters:

Prior to final rendering of the super-resolved images localizations with a lateral uncertainty greater than 150 nm were filtered out. Final images were visualized at 10x magnification (i.e. the image pixel size is 8 nm).

### S6. Selective properties of local gradient algorithm

The Figure S7 shows the selective property of the local gradient algorithms. By varying the window size $r$ one can control the enhancement in the gradient of the corresponding particle.

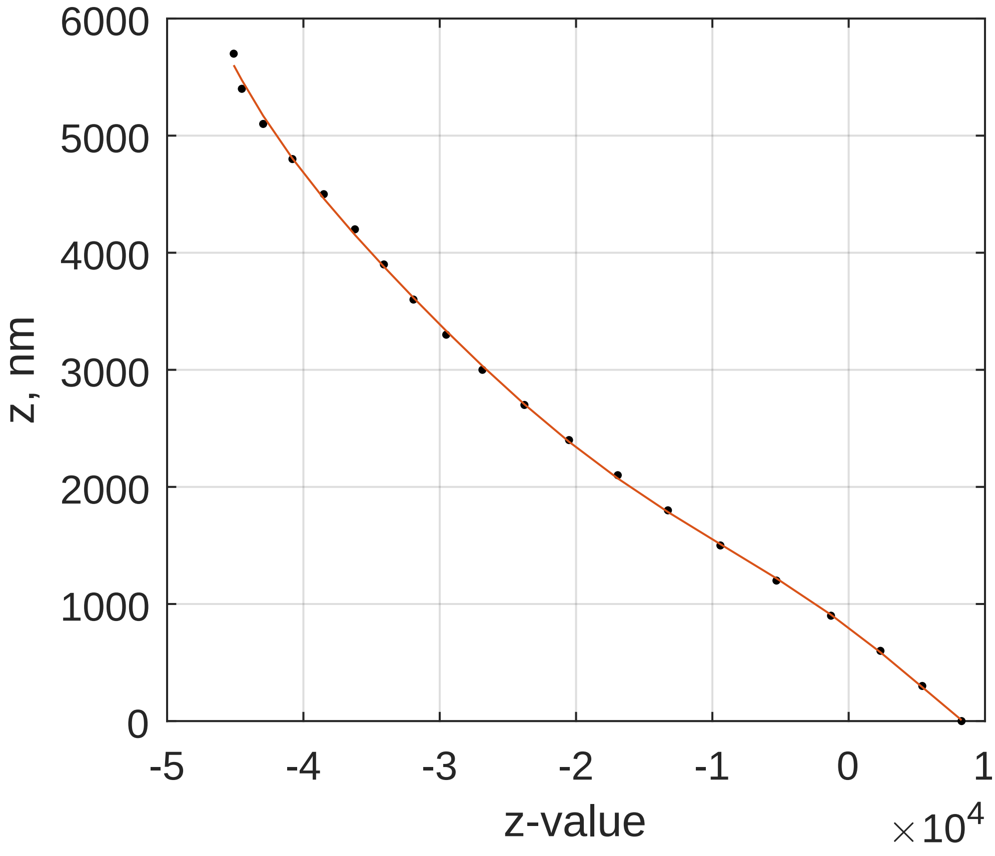


Fig. S6. z-calibration curve for a polystyrene $3\mu m$ particle

| | |
|---|---|
| Image filtering | **Filter**: Gaussian filter<br>**Sigma** [px]: 3.0 |
| Approximate localization of molecules | **Method**: Local maximum<br>**Peak intensity threshold**: 6.5std(Wave.F1)<br>**Connectivity**: 8-neighbourhood |
| Sub-pixel localization of molecules | **Method**: PSF: Elliptical Gaussian (3D astigmatism)<br>**Fitting radius** [px]: 11<br>**Fitting method**: Maximum likelihood<br>**Initial sigma** [px]: 3.0 |
| Visualization of the results | **Method**: Normalized Gaussian<br>**Magnification**: 10<br>**Update frequency** [frames]: 50<br>**3D**<br>**Colorize z-stack**<br>**Z range** (from:step:to) [nm]: -1000:10:1000<br>**Lateral uncertainty** [nm]: 10<br>**Axial uncertainty** [nm]: 20 |

Table 1. Thunderstorm parameters
